# Developmental coupling of cerebral blood flow and fMRI fluctuations in youth

**DOI:** 10.1101/2021.07.28.454179

**Authors:** Erica B. Baller, Alessandra M. Valcarcel, Azeez Adebimpe, Aaron Alexander-Bloch, Zaixu Cui, Ruben C. Gur, Raquel E. Gur, Bart L. Larsen, Kristin A. Linn, Carly M. O’Donnell, Adam R. Pines, Armin Raznahan, David. R. Roalf, Valerie J. Sydnor, Tinashe M. Tapera, M. Dylan Tisdall, Simon Vandekar, Cedric H. Xia, John A. Detre, Russell T. Shinohara, Theodore D. Satterthwaite

## Abstract

To support brain development during youth, the brain must balance energy delivery and consumption. Previous studies in adults have demonstrated high coupling between cerebral blood flow and brain function as measured using functional neuroimaging, but how this relationship evolves over adolescence is unknown. To address this gap, we studied a sample of 831 children and adolescents (478 females, ages 8-22) from the Philadelphia Neurodevelopmental Cohort who were scanned at 3T with both arterial spin labeled (ASL) MRI and resting-state functional MRI (fMRI). Local coupling between cerebral blood flow (CBF, from ASL) and the amplitude of low frequency fluctuations (ALFF, from fMRI) was first quantified using locally weighted regressions on the cortical surface. We then used generalized additive models to evaluate how CBF-ALFF coupling was associated with age, sex, and executive function. Enrichment of effects within canonical functional networks was evaluated using spin-based permutation tests. Our analyses revealed tight CBF-ALFF coupling across the brain. Whole-brain CBF-ALFF coupling decreased with age, largely driven by coupling decreases in the inferior frontal cortex, precuneus, visual cortex, and temporoparietal cortex (*p_fdr_* <0.05). Females had stronger coupling in the frontoparietal network than males (*p_fdr_* <0.05). Better executive function was associated with decreased coupling in the somatomotor network (*p_fdr_* <0.05). Overall, we found that CBF-ALFF coupling evolves in development, differs by sex, and is associated with individual differences in executive function. Future studies will investigate relationships between maturational changes in CBF-ALFF coupling and the presence of psychiatric symptoms in youth.

**SIGNIFICANCE:** The functions of the human brain are metabolically expensive and reliant on coupling between cerebral blood flow and neural activity. Previous neuroimaging studies in adults demonstrate tight physiology-function coupling, but how this coupling evolves over development is unknown. Here, we examine the relationship between blood flow as measured by arterial spin labeling and the amplitude of low frequency fluctuations from resting-state magnetic resonance imaging across a large sample of youth. We demonstrate regionally specific changes in coupling over age and show that variations in coupling are related to biological sex and executive function. Our results highlight the importance of CBF-ALFF coupling throughout development; we discuss its potential as a future target for the study of neuropsychiatric diseases.

## INTRODUCTION

The functions of the human brain are metabolically expensive: despite only weighing 1.5 kg on average, the brain comprises a disproportionate one-fifth of bodily energetic requirements (1–3). To meet such large metabolic demands, the brain receives 20% of cardiac output (4,5). In healthy subjects, the relationship between brain activity and cerebral blood flow (CBF), or neurovascular coupling, is tightly linked at the local level (6–8). Under normal circumstances, metabolites produced during neuronal activity cause vasodilation in the microvasculature, and thus a localized increase in blood flow, to increase glucose and oxygen delivery to active cells (6,9). By coupling metabolic demand (neural activity) and supply (blood flow), the neurovascular unit maintains appropriate energy balance (9).

To measure neurovascular coupling in vivo, proxies for both localized blood flow and regional brain activity are required. Neurovascular coupling has thus been most frequently characterized by relating two neuroimaging-derived measures: CBF and the amplitude of low frequency fluctuations in resting-state blood oxygen level dependent (BOLD) fMRI (ALFF; 10–12). CBF can be measured reliably and without radiation exposure using arterial spin labeled (ASL) MRI (11–13), producing a quantitative measure of blood supply. Although BOLD signal partly reflects changes in CBF, low amplitude fluctuations as measured by ALFF are thought to represent spontaneous brain activity and to underlie intrinsic functional connectivity (14), providing a non-invasive proxy of neuronal function (15,16).

Neurovascular coupling as measured by CBF-ALFF changes in aging, with younger adults displaying higher coupling than healthy older adults (9,10). Sex differences in coupling in adults have also been described (17). Furthermore, decreases in the typically strong coupling between CBF and ALFF have been reported in degenerative and metabolic diseases (9,18,19). Notably, these diseases are associated with significant cognitive impairments, further highlighting the importance of the integrity of this coupling relationship.

Despite this growing literature, notable gaps remain. First, prior studies have examined the relationship between CBF and ALFF primarily at the whole-brain level, yielding a global measure of coupling (20). While informative, such an approach obscures potentially important regional variation in the coupling between CBF and ALFF. Second, previous studies have focused on neurovascular coupling only in healthy and medically ill adults (9,18,19,21). To our knowledge, no prior research has explored how the relationship between blood flow and brain function evolves during childhood, adolescence, and young adulthood. Thus, it is presently unclear how the balance between metabolic demand and supply changes as neurodevelopment unfolds.

Here, we sought to define the local relationship between CBF and ALFF on a regional basis, and to determine how CBF-ALFF coupling evolves over development. To do this, we capitalized on data from the Philadelphia Neurodevelopmental Cohort: a large scale, communitybased study of brain development that included both ASL MRI and fMRI (22,23). We characterized the CBF-ALFF relationship using recently developed tools for inter-modal coupling analysis that allow us to describe the local relationships between these two imaging modalities (24,25). As described below, we extend prior findings demonstrating that blood flow and neural function are strongly related across the brain. Importantly, we also describe how CBF-ALFF coupling evolves in youth, differs by sex, and is related to executive function.

## MATERIALS AND METHODS

### Experimental Design

#### Participants

Participants were drawn from the Philadelphia Neurodevelopmental Cohort (PNC). As previously described, a total of 9,498 participants aged 8-22 years received cognitive assessment and clinical phenotyping, and a subset of 1,601 youths also completed neuroimaging as part of the PNC (22,26). For this report, we excluded participants with missing data, participants currently being treated with psychoactive medication, individuals with medical disorders that could impact brain function, and participants with poor image quality (see below). Eight-hundred thirty-one subjects met criteria and were included in the study (**Figure 1**). The average age of participants was 15.6 years (standard deviation (SD) = 3.4). Forty-two percent (n = 353) were male and 58% were female (n = 478). The institutional review boards of both the University of Pennsylvania and the Children’s Hospital of Philadelphia approved all study procedures.

**Figure 1.**
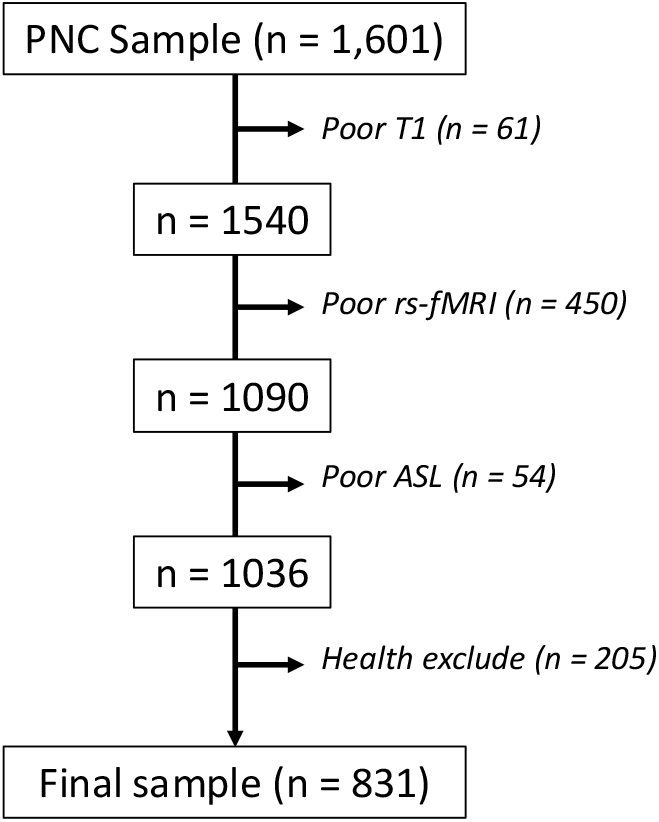
Sample construction. 1,601 participants had neuroimaging scans acquired as part of the PNC. A total of 831 participants were included in the study after excluding participants who failed rigorous quality assessment for poor T1 quality (n = 61), resting state fMRI quality (n = 450), ASL quality (n = 54), and medical and psychiatric comorbidities (n = 205).

#### Cognitive assessment

Cognition was assessed using the University of Pennsylvania Computerized Neurocognitive Battery (CNB) (27). Accuracy of neurocognitive performance was measured across all tests included in the CNB, with raw measures of accuracy being normalized across the entire PNC. Executive function (EF) was summarized using a previously published factor analysis of the CNB (28). Each participant’s *z*-score from this factor analysis was used to evaluate associations between executive function and CBF-ALFF coupling.

#### Image acquisition

PNC imaging was acquired at a single site with a 3T Siemens Tim Trio scanner with a 32-channel head coil (Erlangen, Germany), as previously described (22). To minimize motion, prior to data acquisition, subjects’ heads were stabilized in the head coil using one foam pad over each ear and a third over the top of the head.

High-resolution structural images were acquired to facilitate alignment of individual subject images into a common space. Structural images were acquired using a 3D-encoded magnetization-prepared, rapid-acquisition gradient-echo (MPRAGE) T1-weighted sequence (T_R_= 1810 ms; T_E_ = 3.51 ms; FoV= 180 × 240mm; matrix size =192 x 256, number of slices = 160, slice thickness/gap = 1mm/0; resolution 0.9375 × 0.9375 × 1 mm).

Approximately 6 minutes of task-free functional data were acquired for each subject using a blood oxygen level-dependent (BOLD-weighted) 2D EPI sequence (T_R_ = 3000 ms; T_E_ = 32 ms; FoV = 192 × 192 mm; matrix size = 64×64; number of slices = 46; slice thickness/gap=3mm/0; resolution 3 mm isotropic; 124 volumes). A fixation cross was displayed as images were acquired. Subjects were instructed to stay awake, keep their eyes open, fixate on the displayed crosshair, and remain still. Brain perfusion was imaged with a 3D-encoded spinecho pseudocontinuous arterial spin labeling (pCASL) sequence (T_R_ = 4000 ms; T_E_ = 15 ms; FoV = 220 × 220 mm; matrix size = 96×96; number of slice = 20; slice thickness/gap = 5/1mm; resolution 2.3 x 2.3 x 6 mm; 80 volumes).

### Statistical Analyses

#### Image processing

The structural images were processed using FreeSurfer (version 5.3) to allow for the projection of functional timeseries to the cortical surface (29). Resting-state fMRI scans were processed using a top-performing preprocessing pipeline implemented using the eXtensible Connectivity Pipelines (XCP) (30), which includes tools from FSL (31,32) and AFNI (33). This pipeline included (1) correction for distortions induced by magnetic field inhomogeneity using FSL’s FUGUE utility, (2) removal of the initial 4 volumes for resting-state fMRI, (3) realignment of all volumes to a selected reference volume using FSL’s MCFLIRT, (4) interpolation of intensity outliers in each voxel’s time series using AFNI’s 3dDespike utility, (5) demeaning and removal of any linear or quadratic trends, and (6) co-registration of functional data to the high-resolution structural image using boundary-based registration. Images were de-noised using a 36-parameter confound regression model that has been shown to minimize associations with motion artifact while retaining signals of interest in distinct sub-networks (30). This model included the six framewise estimates of motion, the mean signal extracted from eroded white matter and cerebrospinal fluid compartments, the mean signal extracted from the entire brain, the derivatives of each of these nine parameters, and quadratic terms of each of the nine parameters and their derivatives. Both the BOLD-weighted time series and the artifactual model time series were temporally filtered using a first-order Butterworth filter with a passband between 0.01 and 0.08 Hz to avoid mismatch in the temporal domain (34). The voxel-wise ALFF was computed as the sum over frequency bins in the low-frequency (0.01 - 0.08 Hertz) band of the power spectrum using a Fourier transform of the time-domain signal (14).

As part of image quality assurance, T1-weighted images were excluded for low quality and/or low quality in FreeSurfer reconstruction, as rated by three independent reviewers (35). ASL images were excluded if they had excessive motion (mean relative displacement > 0.5 mm), low temporal signal-to-noise ratio (tSNR < 30), poor image coverage, or an excessive number of voxels that had ceiling intensity values at some point in the timeseries (36,37). Task-free BOLD scans were excluded if the mean relative root mean square (RMS) framewise displacement was higher than 0.2 mm, or if there were more than 20 frames with motion exceeding 0.25 mm (22).

To obtain CBF maps from ASL MRI, images were also processed with tools from XCP (38). CBF was quantified from control-label pairs in the following equation:

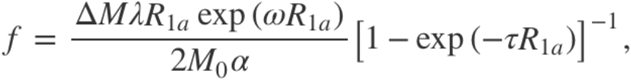

where *f* is CBF, Δ*M* is the difference signal between the control and label acquisitions, *R*_1*a*_ is the longitudinal relaxation rate of blood, *τ* is the labeling time, *ω* is the postlabeling delay time, *α* is the labeling efficiency, *λ* is the blood/tissue water partition coefficient, and *M*_0_ is approximated by the control image intensity (38). We set *α* = 0.85, *λ* = 0.9 g/mL, *τ* = 1.6 s, and *ω* = 1.2 s.

Participant-level CBF images were co-registered to the corresponding T1-weighted image using boundary-based registration with six degrees of freedom (39). Given that T1 relaxation time differs according to age and sex (40-42), the T1 relaxation parameter was modeled on an age- and sex-specific basis (43). This has been shown to enhance the accuracy and reliability of results in developmental samples (36,44).

The CBF and ALFF maps for each individual were projected to the participant’s anatomic surface and smoothed with a 6 mm full-width half-maximum (FWHM) kernel. The smoothed data were normalized to the *fsaverage5* template, which has 10,242 vertices on each hemisphere (18,715 vertices in total after removing the medial wall).

#### CBF-ALFF coupling

Coupling maps were generated at the cortical surface using methods as previously described in detail (24). For each vertex, a 15-mm FWHM neighborhood was defined and a locally weighted regression where ALFF was predicted by CBF was fit. This was repeated at all vertices within each subject, generating one CBF-ALFF coupling map where each vertex was represented by the coupling regression slope (**Figure 2**). A group mean coupling map was also generated by averaging the mean *t*-statistic of each individual’s CBF-ALFF slope at a given vertex.

**Figure 2.**
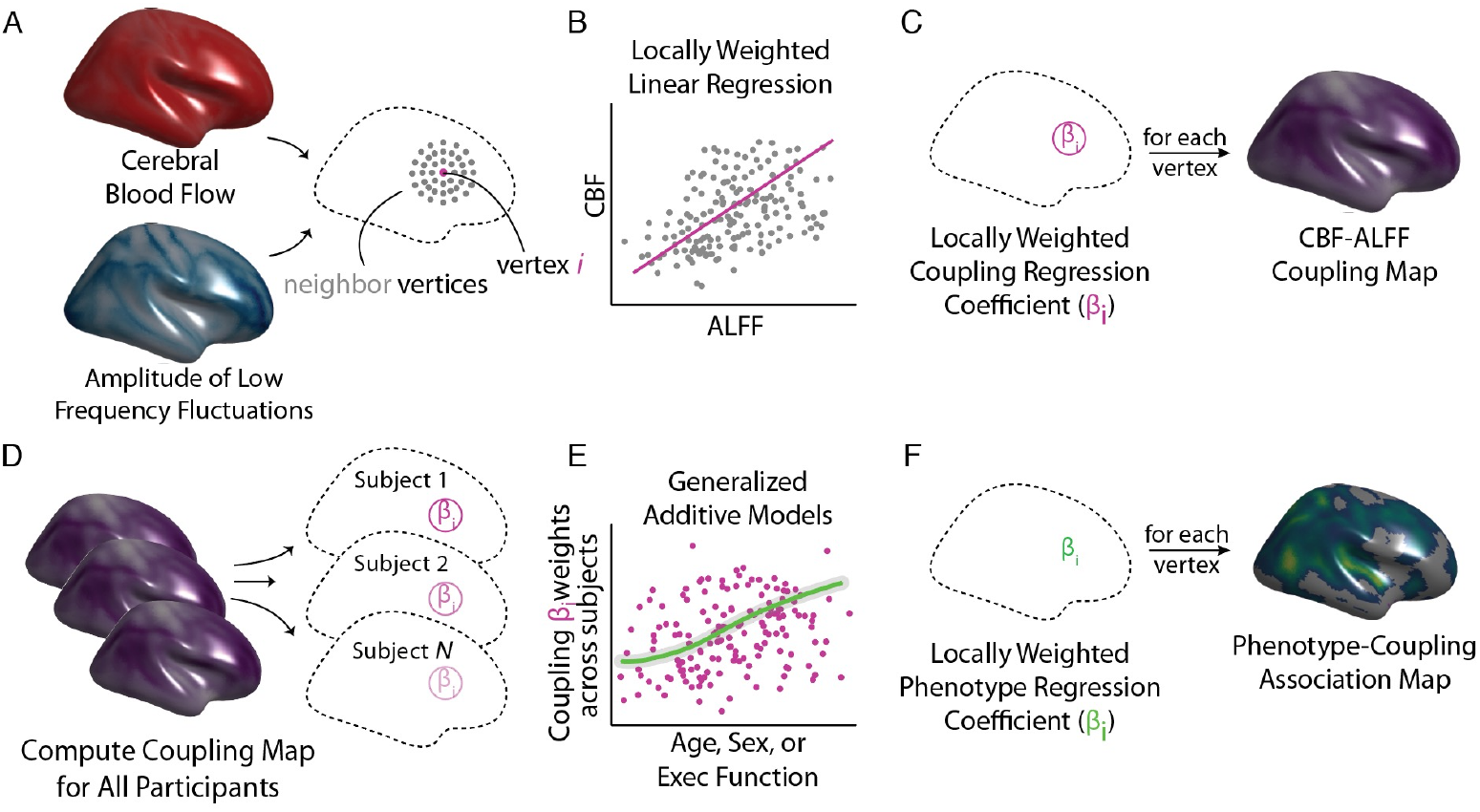
Analysis of CBF-ALFF coupling. CBF-ALFF coupling analysis involves both calculation of within-subject coupling and across-subject comparisons to assess individual differences. **A)** For each subject, a neighborhood for each vertex was identified. **B)** Locally weighted regressions of ALFF onto CBF were calculated. **C)** Locally weighted regressions were repeated at each vertex, resulting in a participant-level coupling map. **D-E)** After subject-level coupling maps were calculated, statistical analyses related covariates of interest (e.g., Age, Sex, and Executive Function) to participant-level coupling maps using generalized additive models (GAMs). **F)** GAMs were fit at each vertex, yielding a group-level statistical map describing individual differences.

#### Statistical analyses and hypothesis testing

We sought to evaluate how neurovascular coupling develops and relates to biological sex and executive function. CBF-ALFF coupling maps for each participant were used for statistical analyses. Generalized additive models (GAMs) were used to calculate linear and nonlinear age and sex effects at each vertex using the following model:

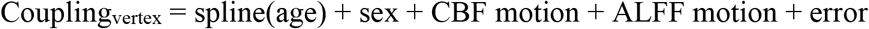

This approach allowed for flexible modeling of both linear and nonlinear effects. For significance testing, smooth terms were fitted as fixed degrees of freedom regression splines (k = 4). We also fit the model with an age-by-sex interaction; however, it was not significant and thus removed from the model. To assess the relationship between coupling and executive function accuracy, we fit a second model which included a linear executive accuracy term in addition to the above model’s covariates (spline of age, sex, CBF and ALFF motion, and error). GAMs were estimated using the **R** package ‘mgcv’ in CRAN (45). All analyses controlled the False Discovery Rate (FDR) at *Q* < 0.05.

In addition to these analyses of effects at each vertex, we also evaluated the relationship between whole-brain coupling and age nonlinearly. We first computed a single mean t-statistic per subject by averaging the t-statistics for the slope at each vertex that met FDR correction, and then used a GAM to estimate the relationship between each subject’s mean coupling and age. To test for windows of significant change across the age range, we calculated the first derivative of the smooth function of age from the GAM model using finite differences, and then generated a simultaneous 95% confidence interval of the derivative using the **R** package ‘gratia’ (45-47). Intervals of significant change were identified as areas where the simultaneous confidence interval of the derivative did not include zero.

#### Network enrichment analyses via spin testing

Given that patterns of neural activity differ across functional networks (48), we sought to characterize whether effects of interest tended to be located within specific functional brain networks (49,50). To do this, we conducted network enrichment analyses. Specifically, we evaluated whether mean coupling and significant associations with variables of interest (age, sex, executive function) were preferentially located in one or more of the seven large-scale functional networks defined by Yeo et al (49). To account for the different size of each network and the spatial autocorrelation of brain maps, statistical testing used a conservative spin-based spatial permutation procedure (51). Briefly, statistical maps from association testing were projected onto a sphere, which was rotated 1,000 times per hemisphere to create a null distribution. For the mean coupling enrichment analysis, the test statistic was the mean *t*-value for each of the seven networks from Yeo et al. For the regression analyses, the test statistic was the proportion of vertices that survived FDR correction. Networks were considered to have significant enrichment if the test statistic in the observed data was in the top 5% of the null distribution derived from permuted data.

#### Code availability

Code for all primary statistical analyses is available at: https://github.com/PennLINC/IntermodalCoupling

## RESULTS

### CBF and ALFF are significantly coupled

Replicating previous findings in adults, we observed robust CBF-ALFF coupling throughout the brain (**Figure 3A**). Coupling was strongest in association cortices, including frontal, parietal, and temporal cortex (*p_fdr_* < 0.05). Network enrichment analysis using spin-based permutation tests revealed enrichment of CBF-ALFF coupling in the default mode network (*p* = 0.045), with nominally higher coupling in the frontoparietal network (*p* = 0.056; **Figure 3B**).

**Figure 3.**
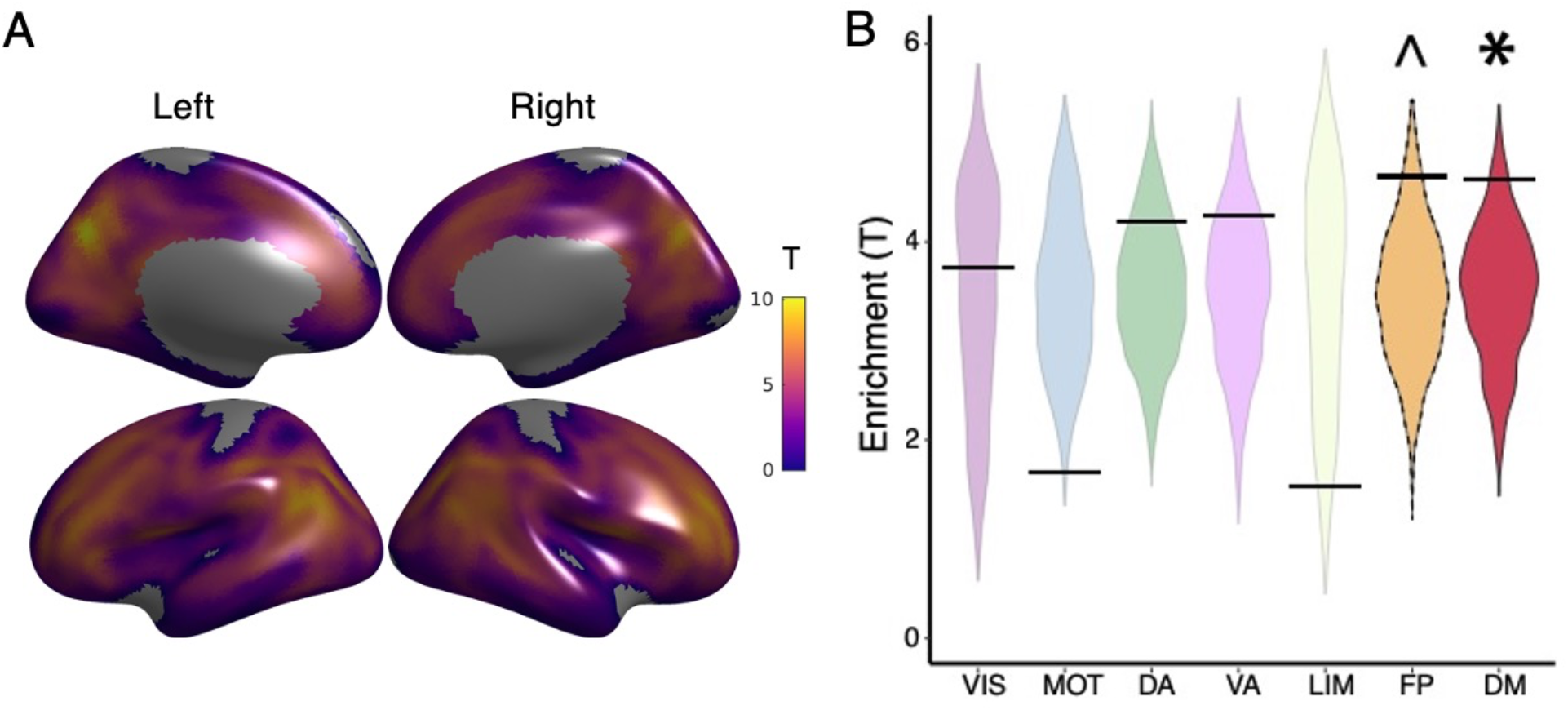
Mean CBF-ALFF coupling. **A)** CBF-ALFF coupling is robust throughout the brain, with maximal coupling in the medial and lateral prefrontal cortex, parietal cortex, posterior cingulate and precuneus. **B)** We used a spin-based spatial permutation test that accounted for spatial autocorrelation to evaluate enrichment of CBF-ALFF coupling in canonical functional networks. This revealed enrichment in the default mode network (*p* = 0.045) with a trend in the frontoparietal network (*p* = 0.056). Star (*) represents statistical significance (*p* < 0.05) and caret (^) represents a non-significant trend (*p* < 0.1). The black bars represent the observed values, whereas the violin plots reflect the null distributions.

### CBF-ALFF coupling declines with adolescent development

Next, we evaluated associations between CBF-ALFF coupling and age. We utilized a generalized additive model to rigorously evaluate both linear and nonlinear developmental effects, while controlling for in-scanner motion and sex. We found that whole-brain mean coupling decreased across development (F_3,828_ = 60.0, *p* < 0.0001; **Figure 4A**). Analysis of the derivative of this spline revealed that coupling significantly decreased between 11.6 years and 20.5 years of age, with a peak decline observed during mid-adolescence at age 16. Fitting this model at each vertex revealed widespread declines in CBF-ALFF coupling across much of the cortex, with peak effects present in the posterior temporal cortex (*p_fdr_* < 0.05; **Figure 4B**). Analyses using spin-based permutation testing revealed enrichment of age-related declines in coupling within the dorsal attention network (*p* = 0.014; **Figure 4C**). In contrast to these spatially extensive declines in coupling, only small regions in bilateral temporal cortices (75 vertices total) showed increased coupling with age.

**Figure 4.**
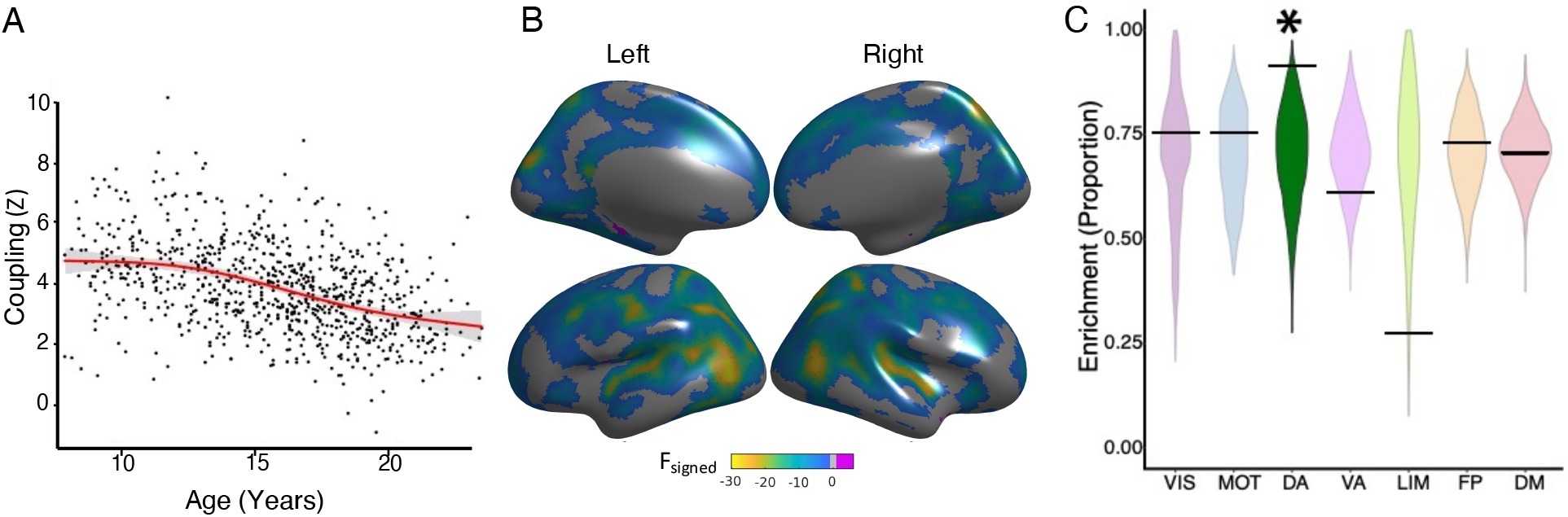
CBF-ALFF coupling evolves with age. Linear and non-linear age effects of CBF-ALFF were flexibly modeled within a generalized additive model at each vertex, while controlling for sex and in-scanner motion; multiple comparisons were controlled using the False Discovery Rate (*Q* < 0.05). **A)** Mean cortical CBF-ALFF coupling declines with age in a nonlinear fashion (F_3,828_ = 60.0, *p* < 0.0001). Data points represent the mean CBF-ALFF coupling (*Z*) for each subject (n = 831) across all vertices that met statistical correction (*p_fdr_* < 0.05). **B)** Vertex-level CBF-ALFF declines were prominent in the posterior temporal cortex, parietal cortex, and dorsolateral prefrontal cortex. For visualization purposes, F_signed_ refers to the F-value from the generalized additive models, with the sign representing directionality (e.g., negative numbers represent lower coupling with age). **C)** Spin testing revealed enrichment of age effects within the dorsal attention network (*p* = 0.014). Star (*) represents statistical significance. Black bars represent the observed values, whereas the violin plots reflect the null distributions.

### CBF-ALFF coupling is higher in females within the frontoparietal network

Having established significant declines in CBF-ALFF coupling with age, we next evaluated sex differences while controlling for both linear and nonlinear effects of age (as well as in-scanner motion). We found that females had stronger coupling than males in bilateral dorsolateral prefrontal cortex, medial frontal cortex, anterior cingulate cortex, and precuneus. In contrast, females had lower coupling in the cuneus and lateral temporal cortex (*p_fdr_* < 0.05; **Figure 5A**). Spin testing revealed that sex differences in coupling were enriched within the frontoparietal network (*p* = 0.034; **Figure 5B**).

**Figure 5.**
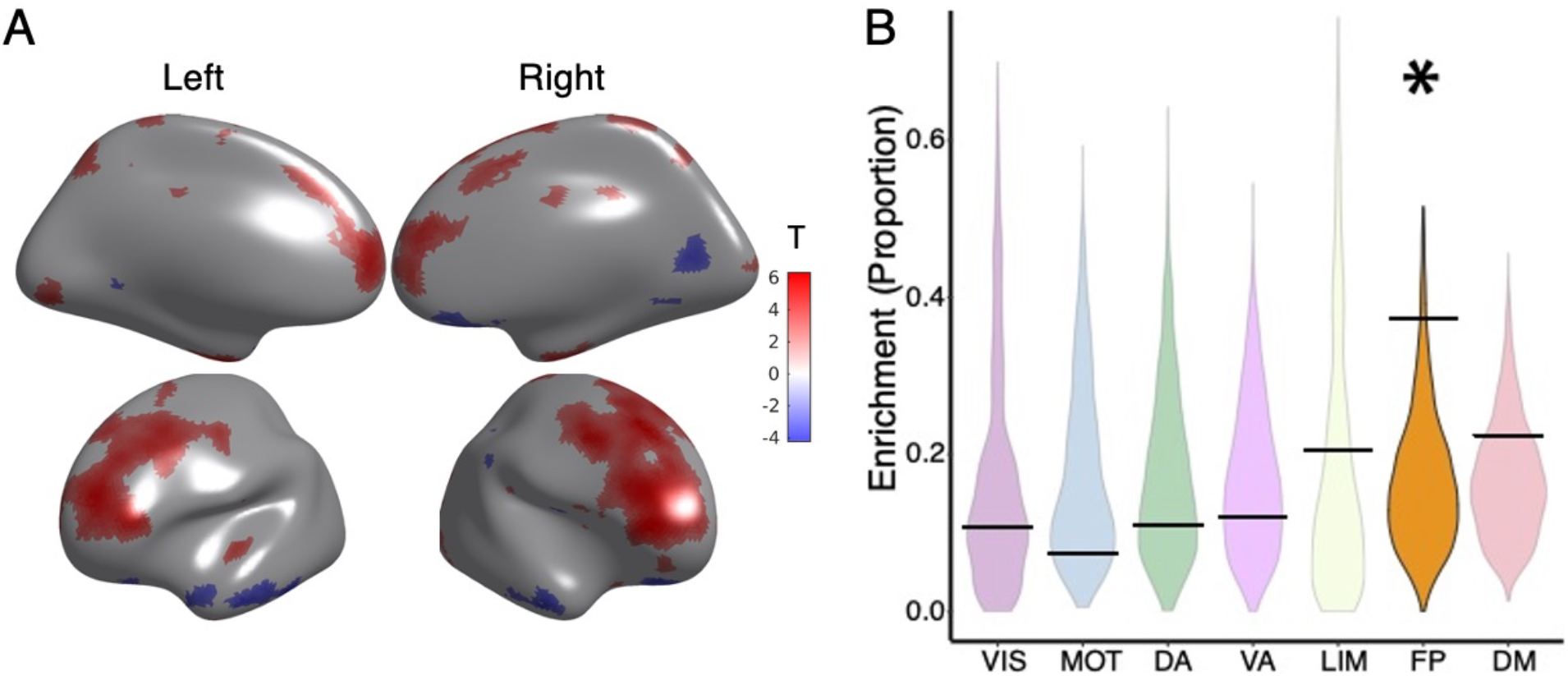
Sex differences in CBF-ALFF coupling. **A)** CBF-ALFF coupling is higher in females than males in the bilateral dorsolateral prefrontal cortex, medial frontal cortex, anterior cingulate cortex, and precuneus. CBF-ALFF coupling differences between females and males were modeled using generalized additive models, while adjusting for both linear and nonlinear age effects as well as in-scanner motion; multiple comparisons were controlled using the False Discovery Rate (*Q* < 0.05). Brain regions where females had greater coupling than males are shown in red, whereas areas where males have greater coupling than females are shown in blue. **B)** Spin testing revealed significant enrichment of sex differences within the frontoparietal network (*p* = 0.034). Star (*) represents statistical significance. Black bars represent the observed values, whereas the violin plots reflect the null distributions.

### CBF-ALFF coupling is associated with executive function

As a final step, we evaluated the relationship between CBF-ALFF coupling and executive function. Throughout, we controlled for linear and nonlinear age effects, sex, and in-scanner motion. We found that better executive function was related to higher coupling in default mode regions including the posterior cingulate cortex, medial prefrontal cortex, and left temporoparietal junction. Furthermore, lower executive function was also related to more coupling in bilateral motor cortex and primary auditory cortex, including Heschl’s gyrus (*p_fdr_* < 0.05; **Figure 6A**). Spin testing revealed enrichment of significant associations with executive function within the somatomotor network (*p* = 0.040; **Figure 6B**).

**Figure 6.**
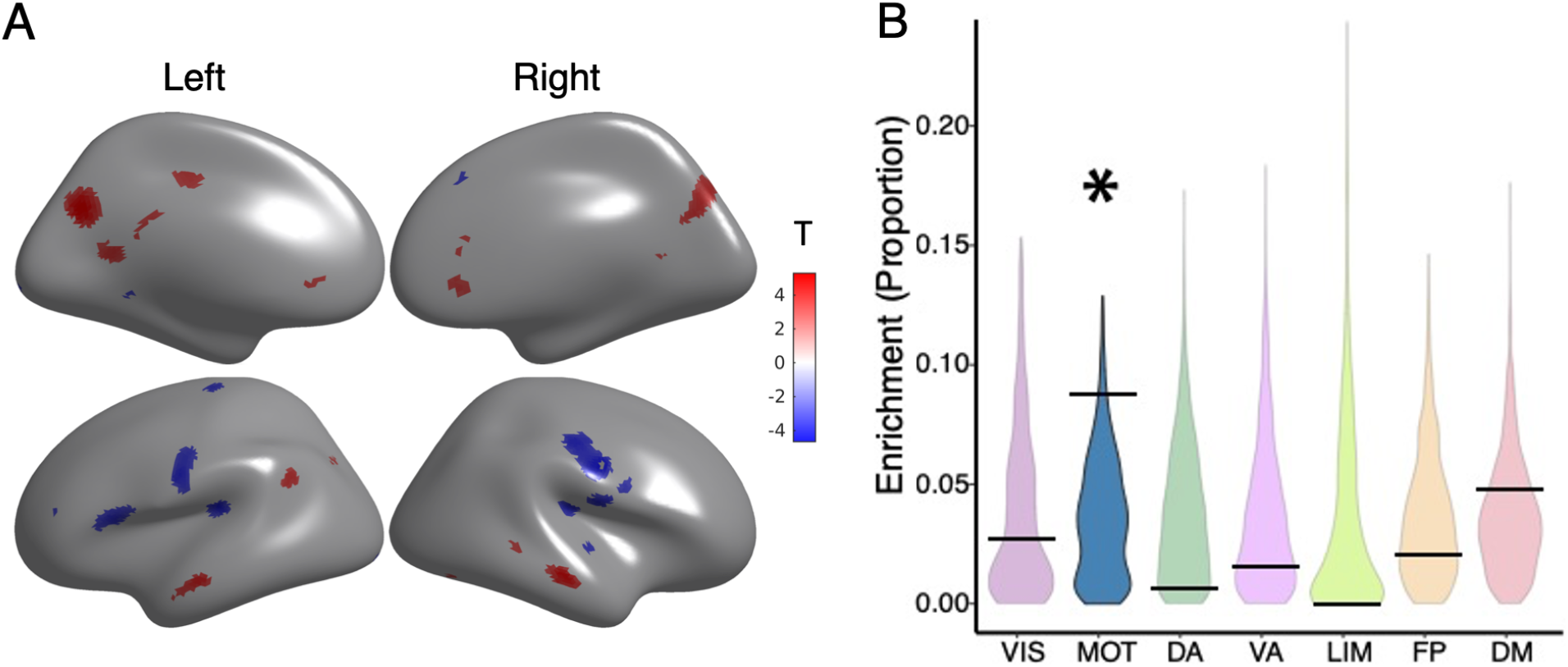
CBF-ALFF coupling is related to executive function. **A)** The relationship of CBF-ALFF coupling to executive function showed regional variation, with both positive and negative associations. Generalized additive models were used to calculate the relationship between CBF-ALFF coupling and executive function while controlling for linear and nonlinear age effects, sex effects, and in-scanner motion; multiple comparisons were accounted for using the False Discovery Rate (*Q* < 0.05). Higher coupling in parts of the default mode were associated with better executive functioning, while higher coupling in parts of the somatomotor network were associated with reduced executive functioning. Brain regions with positive associations are shown in red, whereas areas with negative associations are shown in blue. **B)** Spin testing revealed that associations between executive function and coupling were significantly enriched in the motor network (*p* = 0.040). Star (*) represents statistical significance (*p* < 0.05). Black bars represent the observed values, whereas the violin plots reflect the null distributions.

## DISCUSSION

Using a recently developed statistical method for assessing inter-modal coupling and a large sample of youth, we demonstrated significant CBF-ALFF coupling across the cortex. Furthermore, we identified age-related declines in coupling that were broadly distributed across cortex and were enriched in the dorsal attention network. In addition, we found that sex differences in CBF-ALFF coupling were enriched within the frontoparietal network. Finally, we highlighted the relevance of CBF-ALFF coupling for cognition by showing significant associations between CBF-ALFF with executive function. Taken together, these results extend prior results in adults and demonstrate that CBF-ALFF coupling undergoes a process of developmental calibration that is relevant for cognition.

Neurovascular coupling is thought to reflect the interrelationship between nutrient demand and supply, whereby neuronal activity influences local changes in blood flow (6,7,21,52). At the cellular level, researchers have demonstrated a close relationship between blood flow and neural function that is usually facilitated by communicating astrocytes, where ionic gradients and metabolic biproducts from firing neurons lead to local vasodilation of cerebral arterioles (53,54). As such, high coupling could be conceptualized as an optimized, high fidelity system, where local activity of the vascular unit is influenced by neuronal function (55). Prior work has suggested that CBF-ALFF coupling may be understood as a proxy of neurovascular coupling, allowing a non-invasive window into this process (9,18,19).

Highly metabolically active regions, particularly in association networks that coordinate brain activity across distributed brain regions, tend to have increased CBF as well as ALFF (16,56). Here, we showed that CBF-ALFF coupling is enriched in the default mode network (DMN), with a trend toward significance in the frontoparietal network (FPN). Transmodal association cortices that subserve the DMN and FPN have larger pyramidal neurons with greater spine and synapse density than pyramidal neurons expressed in other parts of the brain (57-59). Similarly, these association networks are the most spatially distributed, with more long-distance cortico-cortical connections (60-63). These neuroanatomical features produce greater metabolic activity and thus may demand a tighter link between activity and blood flow (64).

Though coupling remained tight across development and into young adulthood, CBF-ALFF coupling decreased across much of the cortex, with the greatest rate of change during mid-adolescence. These findings coincide with many previously described structural developmental brain changes (65-67). During the second decade of life, myelin increases while synapses and dendritic spines are pruned to facilitate more efficient between-neuron communication (68–71). To support these structural changes, perineuronal nets surround the cell bodies and dendrites and control ion flow and conduction (72). One possibility is that these structural and ionic adaptations reduce the metabolic demands of the neurons, which has been previously demonstrated in perfusion studies (36,73). These refined local neural circuits may allow for fluctuations in neural activity to be supported by a lower level of metabolic substrate delivered by blood flow.

Sex differences in coupling followed a different pattern than age-related changes. Females showed significantly higher coupling in the FPN, which is essential for cognitive control (74). Interestingly, previous literature has demonstrated that prominent sex differences in perfusion also emerge during the same developmental window, when girls experience an increase in circulating ovarian estrogens (36,37,75). The neuroactive steroid 17B estradiol, the primary estrogen secreted from the ovary, functions as a potent neurovasodilator by enhancing the production of nitric oxide (76). Neurophysiologic studies have consistently demonstrated larger cerebral blood vessel diameter per unit blood pressure in females as compared to males (77,78). Estrogen is also known to selectively increase blood flow in executive areas in adults (79) and emerging evidence has established a link between higher estradiol levels and greater dorsolateral prefrontal cortex activity during emotion regulation in adolescents (80). It is possible that hormone-mediated mechanisms contribute to observed sex differences in coupling within the FPN.

Understanding sex differences in frontoparietal neurobiology that emerge in adolescence is critically important given that mood and anxiety disorders, which are twice as prevalent in girls than boys, also emerge during the same developmental window (81,82). The adult literature consistently reports altered FPN functioning in depression and anxiety (83,84), and these differences can also be identified in development. Previous fMRI research has linked variations in frontoparietal network function with phenotypic heterogeneity in youth depression (85) and anxiety disorders (86). As a next step, it will be important to study whether frontoparietal coupling could be used as a biomarker for sex differences in psychopathology.

The relevance of CBF-ALFF coupling to cognitive function is supported by our results, which detail a significant age-independent relationship between CBF-ALFF coupling and executive function. Specifically, better executive function was associated with lower coupling in the somatomotor network and higher coupling in regions within the DMN, including the posterior cingulate cortex and medial prefrontal cortex. Our findings align with previous unimodal neuroimaging literature on cognition in youth, where executive functioning has been related to both lower-order sensorimotor networks as well as higher-order association networks (87,88). Some previous studies have also suggested a dissociation between how executive functioning relates to neuroimaging measures of sensorimotor and association networks (89). Our findings may further indicate that the development of executive function in youth relies on a balance between decreased coupling in lower-order networks, and increased coupling in higher-order association networks. It is also possible that this dissociation is specific to the adolescent developmental stage, where lower coupling in somatomotor network indicates a refined circuit, whereas higher correlation in the DMN indicates that the system is undergoing developmental tuning and would presumably move to a lower coupling state in adulthood.

There are several limitations to our study which should be noted. Previous authors have understood CBF-ALFF coupling as a representation of neurovascular coupling, which has framed our understanding of this measure (12,18,19). However, we are not able to measure neurovascular coupling directly and had to rely on proxy measures to characterize neurovascular coupling *in vivo*. Typically, positron emission tomography has generally been the gold standard for measuring blood flow, whereas we use ASL to quantify CBF. However, previous studies have demonstrated good correlation between PET and ASL (13). Given the potential risks of radiologic exposure to children with PET scanning, CBF as measured by ASL is a much safer method for use in large-scale studies of youth (90–92). Additionally it should be noted that ALFF is derived from the BOLD signal, which inherently has a vascular component (93). However, numerous past studies have suggested that ALFF signals reflect neural activity (14,15,94,95). Lastly, we evaluated a cross-sectional sample which prevents us from estimating within-individual change; future studies in longitudinal samples will be important.

The limitations notwithstanding, we provide new evidence for the developmental evolution of CBF-ALFF coupling in youth, as well as distinct associations with sex and executive function. Our findings suggest numerous avenues for future study. Longitudinal assessments that allow for the measure of within-subject change in coupling may help us better understand normal physiologic brain development at a personalized level. Additionally, transdiagnostic deficits in executive functioning are commonly observed across psychiatric illnesses (96,97). Given that deficits in executive functioning were associated with altered CBF-ALFF coupling, future studies could yield important insights into whether regional differences in CBF-ALFF coupling are linked to the onset and development of psychopathology. Eventually, characterizing neurovascular coupling in youth at risk may aid in the development of targeted pharmacologic and neurotherapeutic treatments.

## Acknowledgements

This work was supported by grants from the National Institute of Mental Health (NIMH; Grant Numbers: R01MH112847 to TDS and RTS; R01MH120482, and R01MH113550 to TDS; R01 MH123550-01 to RTS; R01MH107235 to RCG; 2T32MH019112-29A1 to EBB; T32MH014654 to BLL; F31 MH123063-01A1 to ARP; R01MH120174, R01MH119185, and R56AG066656 to DRR; DGE-1845298 to VJS. The PNC was funded by RC2 grants MH089983 and MH089924 to REG from the NIMH.

